# Theta phase and theta-gamma coupling organise the spoken language network

**DOI:** 10.64898/2026.06.03.729880

**Authors:** Haya Akkad, Daniel Bush, Robert Seymour, Tae Twomey, Susanne Pelke, Sasha Ondobaka, Sven Bestmann, Jenny Crinion

## Abstract

Speech production requires rapid coordination of conceptual and lexical processes across distributed cortical networks, yet the neurophysiological mechanisms enabling this coordination remain poorly understood. Oscillatory coupling has emerged as a candidate mechanism for coordinating neural activity across spatial scales. Here, we used whole-head magnetoencephalography during overt picture naming to test how phase and phase-amplitude coupling organise neural dynamics preceding articulation. We show that theta (4-8 Hz) phase coupling increases within two functionally distinct networks: a ventral occipito-temporal network supporting object recognition and a medial fronto-temporal network supporting semantic-lexical retrieval. These networks converged in the right fusiform gyrus beyond chance levels, identifying a candidate integration hub. In parallel, whole-brain analysis of theta-gamma (4-8 Hz, 40-100 Hz) phase-amplitude coupling revealed selective increases in the left inferior frontal and fusiform gyri during picture naming relative to control. Mixed-effects modelling further showed that coupling in the left fusiform correlates with trial-level response times during naming but not control trials. Together, these findings reveal the oscillatory mechanisms that implement known functional specialisation in the spoken language network; theta phase coupling coordinates distributed recognition and retrieval streams, while theta–gamma coupling modulates local computations within core word production nodes. By defining an oscillatory framework for real-time speech production, this work advances mechanistic understanding of the spoken language network and identifies frequency- and region-specific targets for neuromodulation of language production disorders.

**Highlights:**

- Distinct oscillatory mechanisms support distributed and local language production processes
- Theta phase-coupling distinguishes object recognition and word retrieval networks
- Theta-gamma coupling increases locally in core language regions
- Increases in theta-gamma coupling scale with trial-by-trial naming speed

## Introduction

Language production is a fundamental aspect of human behaviour, and its disruption is one of the most common ^1,2^ and feared outcomes following brain injury and disease ^3,4^. Despite this, the neurophysiological mechanisms underlying the spoken language network are poorly understood, limiting the efficacy of neuromodulation for language production disorders ^5-7^.

At the neural level, language production requires rapid coordination of conceptual, semantic and lexical processes across distributed cortical networks ^8,9^. These processes involve at least two major functional streams; ventral occipito-temporal systems supporting object recognition and medial fronto-temporal systems supporting semantic-lexical retrieval ^8,10,11^ (Box 1). However, the neural mechanisms supporting their large-scale coordination remain unclear. Neural oscillations provide a candidate mechanism: low frequency phase-coupling coordinates activity across distributed networks ^12-14^ and can interact with higher frequency activity within a given region via phase-amplitude coupling (PAC) ^12,15^. These principles have been identified across subcortical and neocortical circuits, where they are increasingly implicated in coordinating behaviour across sensory, motor and cognitive domains ^16-20^.

Theta activity is particularly relevant to language production. Theta power increases are consistently reported during semantic and lexical processing in language production tasks ^21-24^. Additionally, long-range theta phase coupling predicts developmental verbal ability, more than alpha or beta rhythms ^25^. Theta-gamma PAC is increasingly recognised as a conserved cortical mechanism for coordinating local neural activity within large-scale networks ^12^. However, while theta dynamics have been associated with spoken language, their role in coordinating distributed processing during language production remains unclear.

Here, we hypothesised that complementary theta and gamma coupling dynamics organise the spoken language network across spatial scales. Using whole-head magnetoencephalography during overt picture naming, we quantified theta phase coupling and theta-gamma PAC preceding articulation. Our findings define an oscillatory framework for real-time coordination of distributed recognition and retrieval streams in the spoken language network and identify mechanistic targets for neuromodulation of language production disorders.

## Results and Discussion

### Behavioural paradigm and performance

Twenty-seven healthy older adults (9 female, mean age 64 ±6.89 years) completed an overt picture naming task including a matched control condition. To enhance translational relevance, participants were age-matched to clinical cohorts with language impairment. The naming condition consisted of line drawings of everyday objects; the control condition used scrambled versions of the same objects preserving low-level visual information. Participants were instructed to name the object as quickly and accurately as possible or respond ‘no’ if the image was visual noise, ensuring motor output in both conditions (**Figure 1A**). Participants named objects more slowly than they rejected scrambled images (t = 13.9; P < 0.001). The mean response time (RT) measured from picture onset was 1.08s (SD: 0.11) in the naming condition and 0.81s (SD: 0.11) in the control condition (**Figure 1B**). Invalid responses, including dysfluencies and multiple or missing responses, were coded as ‘missed’ trials and excluded from all analyses. Misses were rare: 2.93% (SD: 3.15) in the naming and 0.69% (SD: 1.33) in control.

**Figure 1.**
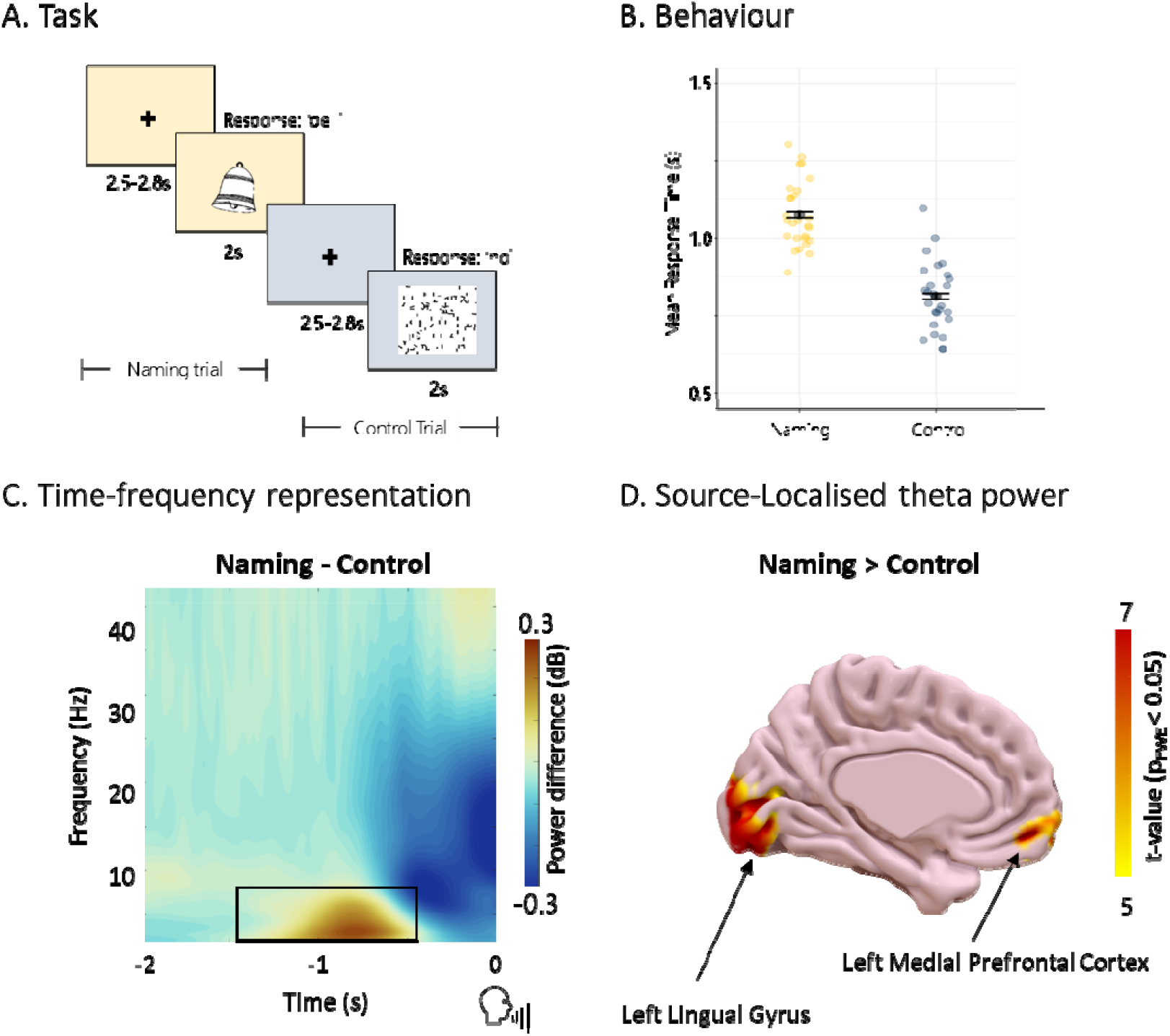
Experimental design and behaviour. **A)** Task. Each trial began with a fixation cross centred on the screen for a jittered period between 2.5 and 2.8 s, followed by the stimulus for 2 s. Stimuli consisted of 200 line-drawings of common objects, displayed unaltered for the naming condition and scrambled for the control condition (400 total trials). Stimuli were pseudorandomised, ensuring no condition appeared more than four times in a row. Participants were instructed to name the object as quickly and accurately as possible or respond ‘no’ if the image was visual noise. Trials were divided into 4 blocks of 100 trials each, with self-paced breaks in between. **(B)** Mean response times (RT) for the naming (yellow, mean RT = 1.08 s) and control (blue, mean RT = 0.8 s,) condition. Responses were significantly faster in the control condition (P < 0.001). Error bars demonstrate standard error of the mean (SEM). **(C)** Time-frequency representation (TFR) of power time-locked to response onset. TFR of the difference in relative power change from baseline (200ms pre-stimulus onset) during naming relative to the control condition. Black outline represents the spectrotemporal window of interest used for source reconstruction (4–8Hz, -1.4 to -0.6s). **(D)** Source of theta power during naming relative to control. Peak regions with the strongest theta (4-8 Hz) power increases were in left lingual gyrus [MNI: -14 -96 -12] and left medial prefrontal cortex [MNI: -8 56 -10]. Results are displayed at a voxel-level threshold of P < 0.05

### Theta phase coupling coordinates distributed processing streams

Prior studies have highlighted the relevance of theta activity for spoken language, but its mechanistic role remains unclear. Theta phase coupling between frontal and occipito-temporal EEG sensors correlates with lexical demand in adults^26^ and predicts developmental verbal ability in children and adolescents^25^. However, how theta phase coupling organises the spoken language network is poorly understood.

To address this, we quantified source-resolved theta activity across the whole brain while participants completed the picture naming task, focusing on the 1 s preceding articulation. Contrasting theta power between naming and control identified two whole-brain-corrected clusters in the left posterior lingual gyrus and left medial prefrontal cortex (mPFC) (**Figure 1C, D, Supplementary Table 1**). Examining theta phase coupling between these regions and the rest of the brain revealed dissociable fronto-temporo-occipital processing streams supporting object recognition and semantic-lexical retrieval (**Figure 2)**.

**Figure 2.**
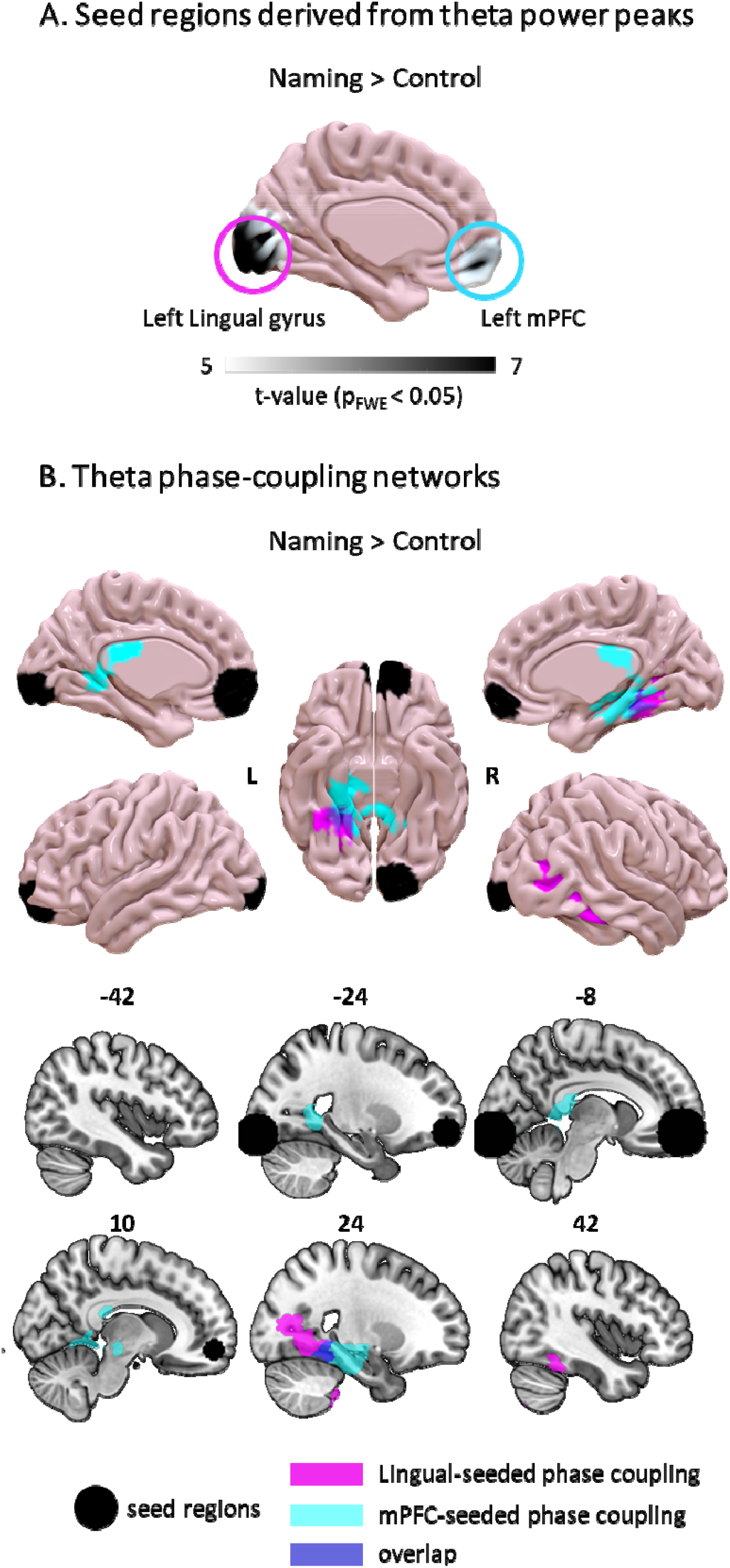
Theta (4-8 Hz) phase coupling in distinct language production networks. **(A)** Seed regions were defined based on voxels showing the greatest increase in theta power during picture naming relative to the control condition. **(B)** Seeds are shown in black and represent a 20mm radius centred on a voxel in the left lingual gyrus [MNI(-14 -96 -12)] and the left medial prefrontal cortex (mPFC) [MNI(-8 56 -10)]. Theta phase coupling was estimated between each seed and all other voxels across the whole brain and summary images were contrasted between conditions. Resulting networks are overlayed on the cortical surface (top) and orthogonal sections (bottom). The left lingual seed showed theta phase coupling with a right ventral occipito-temporal network (cyan) associated with object recognition. The left medial prefrontal seed displayed theta phase coupling with bilateral medial temporal regions (magenta) consistent with semantic-lexical retrieval networks. The right fusiform gyrus emerged as a region of overlap between the two networks (purple). The extent of spatial overlap in the fusiform significancy exceeded chance levels (p < 0.001; see Supplementary Figure 2). Results are displayed at a voxel-level threshold of *P* < 0.001 and a cluster-level family-wise error–corrected threshold of P_FWE_ < 0.05 for naming relative to control trials. See supplementary table 2 for corresponding MNI co-ordinates.

Specifically, posterior theta (4-8 Hz) activity in the left lingual gyrus showed increased phase-coupling with a right ventral occipito-temporal network during naming compared to control, including right lingual and fusiform gyri (**Figure 2A**, see Supplementary Table 2A for peak MNI coordinates). Regions in this network fell within the ventral visual processing stream, known to be crucial for object recognition and semantic knowledge ^27-30^, consistent with dual-stream models of cortical organisation^31,32^. Coupling with the right hemisphere, particularly the right fusiform, is consistent with this region’s role in visually guided word retrieval^33-35^.

In contrast, theta in the left medial pre-frontal cortex (mPFC) was phase-coupled with bilateral regions in the medial temporal lobe (mTL), including the fusiform, hippocampal and parahippocampal cortex (**Figure 2B**, see Supplementary Table 2B for peak MNI coordinates). mPFC-mTL interactions, including theta phase coupling, are linked to long-term memory retrieval ^36,37^. Notably, the fusiform and parahippocampal regions identified in this network are consistently associated with semantic-lexical retrieval during word production^8,32,35,38-40^, and their disruption via direct stimulation or lesion results in semantic and lexical retrieval deficits during naming ^40-43^.

Finally, the two networks converged spatially in the right fusiform gyrus, identifying a candidate integration hub linking recognition and retrieval processes (**Figure 2)**. This is consistent with fMRI evidence linking right fusiform activation specifically to picture naming^33,35^, a task requiring integration of visual object recognition and lexical-semantic retrieval, relative to word retrieval cued by sentence context or definition. Permutation testing confirmed that this overlap exceeded chance levels (p < 0.001; Supplementary Figure 1).

Together, these findings show that distinct recognition and retrieval processing streams are coordinated via theta phase coupling during language production, converging in the right fusiform. This extends dual-stream models of cortical processing^31,32^ to speech production and grounds large-scale spoken language organisation in established oscillatory principles of neural integration^12-14^.

### Theta-gamma coupling modulates local computations

In addition to large-scale theta phase coupling, we tested our hypothesis that theta-gamma phase-amplitude coupling (PAC) supports local computations during language production. Intracranial recordings from frontal and temporal electrodes show theta-gamma PAC occurs during a wide range of tasks and its spatial distribution varies in a task-relevant manner^16^. However, the regional organisation and behavioural relevance of PAC during language production is unknown.

Here, we quantified theta-gamma PAC across the whole brain using source-resolved MEG analyses during the picture naming task, contrasting naming and control in the 1 s preceding response. This identified two regions that survived multiple-comparison correction across the whole brain: the pars triangularis of the left inferior frontal gyrus (LIFG) (MNI [-38; 28; 2]) and the left middle fusiform gyrus (MNI [-28; -10; -30]) (**Figure 3A**, Supplementary Table 3). Both regions are established core nodes in the word production network^35,40,43^. Next, we used a comodulogram analysis ^16^ (see Methods) to test whether PAC in these regions was driven by specific theta or gamma sub-bands during naming compared to control. This revealed that significant PAC during naming occurred between 5-7 Hz theta and 58-78 Hz gamma in the LIFG, while coupling in the left fusiform occurred over a broader range between 4-8 Hz theta and 42-92 Hz gamma (**Figure 3B**). PAC was frequency and region specific; we found no evidence for alpha-gamma or beta-gamma coupling during naming compared to control across the whole brain. Additionally, coupling remained significant after regressing colinear changes in theta and gamma power, confirming results were not driven by concomitant power increases that can affect the signal-to-noise ratio.

**Figure 3.**
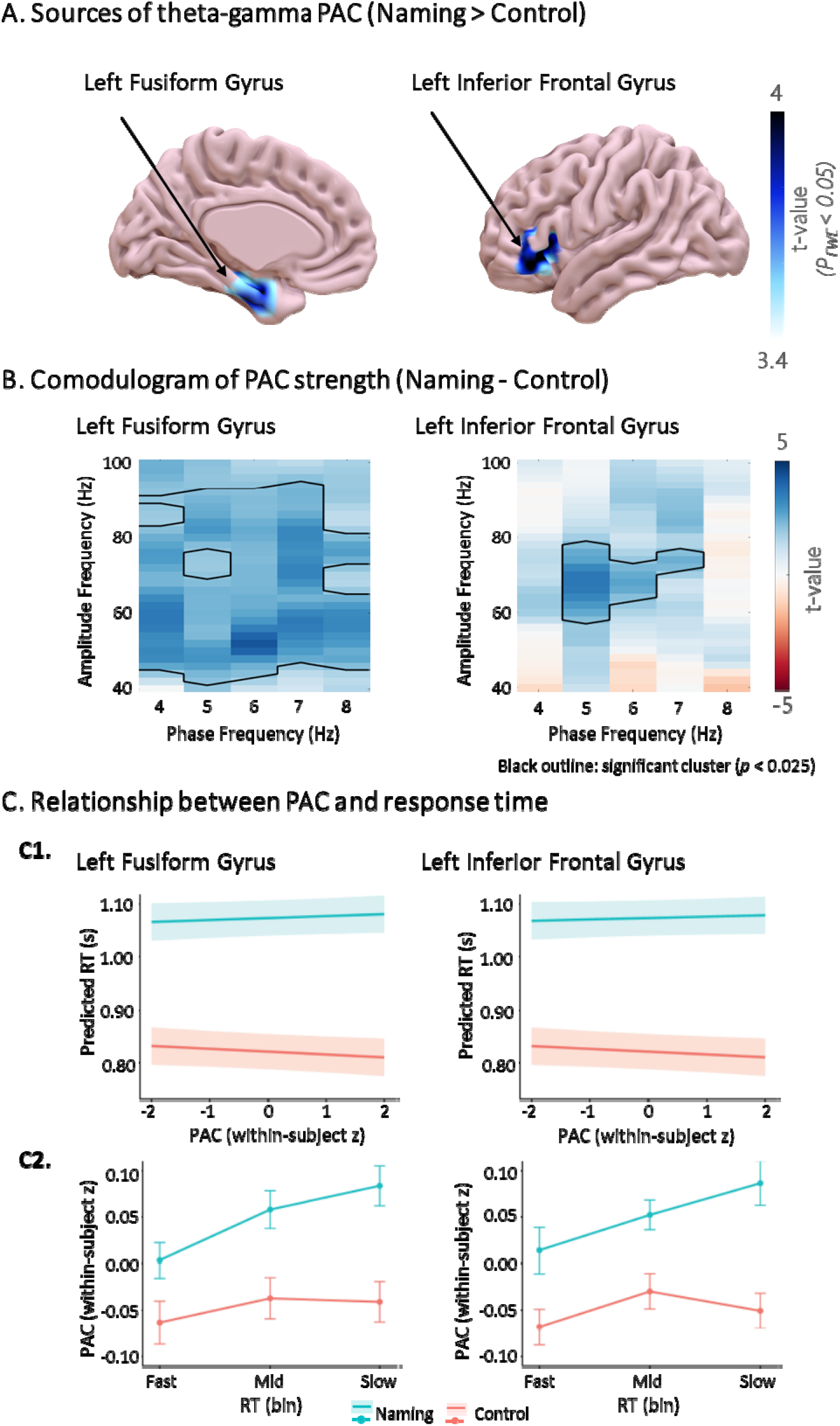
Theta-gamma phase-amplitude coupling increases in core language production regions during naming. **A)** Theta (4–8 Hz)-gamma (40–100 Hz) phase-amplitude coupling (PAC) was estimated at each voxel across the whole-brain volume, in the 1 s preceding response onset. Summary images were compared between naming and control trials using a general linear model in SPM12 (voxel-level threshold P < 0.001; cluster-level P_FWE_ < 0.05). Significant increases in theta-gamma PAC during naming relative to control were found in the left middle fusiform gyrus (left) and the left inferior frontal gyrus (LIFG; right). **B)** Phase-to-amplitude comodulograms contrasting naming and control conditions, computed from source-reconstructed timeseries from the left fusiform gyrus [MNI: -28; -10; -30] and LIFG [MNI: -38; 28; 2]. Black outlines indicate statistically significant clusters (P < 0.025, cluster statistics). In the left fusiform gyrus, coupling spanned a broad theta-gamma range (4-8 Hz theta phase; 42-92 Hz gamma amplitude; peak: 6 Hz / 52 Hz), whereas LIFG coupling occurred in a narrower frequency range between 5-7 Hz theta phase and 58-78 Hz gamma amplitude (peak 5Hz / 68Hz) **C)** Theta-gamma PAC in the left fusiform gyrus showed a significant condition-dependent relationship with response times (RTs) (p = 0.039). The corresponding interaction in the left inferior frontal gyrus (LIFG) did not reach significance (p = 0.07). **C1)** Shows predicted RT as a function of trial-by-trial PAC in the left fusiform gyrus and LIFG, estimated from linear mixed-effects models (random intercept for subject). PAC values were z-scored within subject to capture trial-to-trial fluctuations. Lines represent model-predicted RT for each condition, and shaded regions indicate 95% confidence intervals. **C2)** Shows the same relationship visualised by binning trials into subject-wise RT tertiles (fast, mid, slow) for display purposes only. Points represent the mean PAC across subjects within each bin and condition, with error bars showing ±1 SE across subjects. Statistical inference was performed on trial-level data using the mixed-effects models.

### Theta-gamma coupling predicts response time

Finally, we asked how theta-gamma PAC within these regions relates to behaviour. Using trial-level linear mixed-effects models with subjects as a random intercept, we found that in the left fusiform, stronger PAC was associated with slower responses during naming but not control trials, yielding a significant PAC x condition interaction (β = 0.0090, SE = 0.0044, t(10150) = 2.04, p = 0.041). This effect remained significant when including subject-specific random slopes for condition (F(1, 10121) = 4.27, p = 0.039), indicating robustness to inter-individual variability in task effects. A similar pattern was observed in the LIFG but did not reach significance (β = 0.0079, SE = 0.0044, t = 1.77, p = 0.076) **(Figure 3C1, 3C2)**.

These findings suggest that theta-gamma PAC indexes the engagement of local processing within core naming regions. Coupling in these regions was specific to naming behaviour, relative to control, and stronger coupling accompanied slower responses, consistent with greater local processing demands.

Converging evidence from fMRI, electrocorticography and direct stimulation using depth electrodes consistently identify the left fusiform gyrus and left inferior frontal gyrus in core processes of word production, independent of input modality ^35,40,43^. Our findings identify theta-gamma PAC as a candidate mechanism modulating local processing within these key nodes.

### Conclusion and Significance

Speech production is coordinated by frequency-specific oscillatory mechanisms operating across spatial scales. Theta phase coupling coordinates distinct object recognition and word retrieval streams that converge in the right fusiform gyrus, while theta-gamma PAC increases within core word production nodes in the left inferior frontal and fusiform gyri and scales with trial-by-trial naming speed.

This oscillatory framework maps onto known bilateral fusiform specialisation. fMRI studies of word production show that that the right fusiform preferentially supports *visually* guided word retrieval^33,35^, whereas the left fusiform supports modality-independent lexical-semantic processing^35,40,43^. Our findings reveal that these lateralised fusiform functions are implemented via complementary oscillatory mechanisms, with theta phase coupling coordinating integration of visual recognition and word retrieval streams in the right fusiform, and theta-gamma PAC supporting local computations within the modality-agnostic left fusiform. Together, the present results define an oscillatory framework for real-time language production, advancing mechanistic understanding of the spoken language network and identifying targets for causal and translational investigation.

Clinically, these findings have direct translational relevance for anomia, a severe word retrieval deficit^44^, and related language production disorders. Current neuromodulation approaches remain largely nonspecific, resulting in variable therapeutic outcomes^45^. Our findings provide frequency- and region-specific targets for future mechanistically grounded neuromodulation^20^.

### Limitations and considerations

Inferences regarding specific linguistic processes are necessarily limited by the use of a low-level control condition designed to capture all pre-articulatory stages during object naming. Additionally, limiting stimuli to monosyllabic, high-agreement items ensured comparable motor preparation and response latencies between conditions and facilitates future replication and comparison in clinical populations with impaired language ability. However, future studies could extend the present findings by examining the mechanisms identified here under varying retrieval, selection, and phonological demands. Finally, the causal contribution of theta phase and theta-gamma coupling to naming remains to be determined. Future studies may consider using brain stimulation to test how engagement or disruption of these coupling dynamics alters naming behaviour in healthy individuals ^20^ and how they relate to language deficits in clinical populations.

## Acknowledgements

This work was supported by Brain Research UK – BRUK Doctoral Fellowship (H.A) and the Wellcome – 203147/Z/16/Z and106161/Z/14/Z (J.C). The funders had no participation in the design and results of this study.

## Declaration of interests

The authors declare no competing interests.

## Materials and Methods

### Experimental model and subject details

Twenty-seven healthy right-handed, older adults (9 female, age 64 ± 6.89, mean ± SD) participated in the experiment. Participant selection was based on the following inclusion criteria: 1) Fifty+ years of age (matched to stroke survivors), 2) proficient in spoken and written English. All participants had normal or corrected-to-normal vision and no history of neurological or language deficits.

The study was conducted according to the Declaration of Helsinki and was approved by the University College London research ethics committee. All participants provided written informed consent.

### Method details

#### Data collection

MEG recordings were made using a 275-channel axial gradiometer system (CTF Omega, VSM MedTech) sampling at 600 Hz while participants sat upright in a magnetically shielded room. Head position coils were attached to nasion and left and right pre-auricular sites for anatomical co-registration. Eye movements were recorded using an Eyelink 1000 eye tracker (SR Research). Stimulus presentation was controlled using the Psychophysics Toolbox v.3 (Psychtoolbox-3) in Matlab (2022a).

#### Experimental design

A list of 200 target words for common objects was drawn from the International Picture Naming Project (IPNP; http://crl.ucsd.edu/experiments/ipnp/index.html; Szekely et al., 2004) and the MRC Psycholinguistic Database (http://websites.psychology.uwa.edu.au/school/MRCDatabase/mrc2.html; Coltheart, 1981). All object names were monosyllabic with high naming-agreement. The materials are listed in supplementary Table 4. For the naming condition, 200 line-drawings of common objects were derived partly from the IPNP, and the remainder found on the internet (with similar style/figurative features as the IPNP items). For the control condition, the same 200 images were scrambled so as to become unrecognisable whilst maintaining the same amount of low-level visual information. The stimuli were pseudorandomised, ensuring that no condition appeared more than four times in a row. Participants were instructed to name the object as quickly and accurately as possible or verbally respond ‘no’ if the image was visual noise – this ensured the control condition involved motor output, but no semantic or lexical retrieval.

Before starting the experiment, participants completed a short practice with 8 trials both inside and outside the scanner, to familiarise them with the task. Stimulus items used in the practice session were not included in the experiment. A trial began with a fixation cross centred on the screen for a jittered period between 2.5 and 2.8 s, followed by the stimulus for 2 s. The trials were divided into 4 blocks of 100 trials each, with self-paced breaks in between (**Figure 1A**).

#### Data preprocessing and artefact rejection

The data were imported into SPM12^46^ (Wellcome Centre for Human Neuroimaging, London, UK) and downsampled to 300 Hz. Eye blink and heartbeat artefacts were identified by independent component analysis (ICA) and removed using FieldTrip^47^ and EEGLAB^48^. High-pass (0.5 Hz) and notch (48-52 Hz) filters were applied and the data were epoched and merged across sessions. Trials containing signal artefacts (3.93% of all trials) were identified and removed using a Generalized Extreme Studentized Deviate (GESD) test to find outliers, with a threshold of α=0.05, from the OHBA Software Library (OSL; https://github.com/OHBA-analysis/osl) adapted for Matlab (2022a). Additionally, as our analyses were response-locked, trials where no response time was detected - either due to a missed trial (no-response, delayed response or dysfluencies) or a technical fault in the MEG audio channel – were identified and removed by visual inspection (on average, 2.93% naming trials and 0.69% control trials). This resulted in an average of 194.07 [standard deviation (SD = 6.31)] naming trials and 196.52 (SD = 2.65) control trials, per subject. Data was segmented using either the stimulus onset or response time of each individual trial. Response time was identified from the MEG audio channel. Stimulus onset was detected by placing a photodiode on the display screen in the magnetically shielded room; this produced a change in voltage when the stimulus appeared and was recorded alongside the MEG signals.

### Quantification and statistical analysis

#### RT analysis

Vocal responses were recorded on a trial-by-trial basis using a microphone in the magnetically shielded room and evaluated offline. Responses containing disfluencies, delayed or no response were coded as invalid and their corresponding trials were excluded from all analyses. Average RTs across participants were compared in each condition (naming versus control) using a paired samples t-test.

### MEG data analysis

#### Theta Power

##### Response-locked Time-Frequency Analysis

Estimates of dynamic oscillatory power were obtained by convolving the MEG signal with a five-cycle Morlet wavelet. Power values were obtained for 30 logarithmically spaced frequency bands in the 2–100 Hz range, from 4 s before response onset to 2 s after response onset in each trial (i.e. centred on the 2 s period preceding a response, with 2 s padding on either side). To obtain measures of power change in each frequency band from baseline, log-transformed power values at each time point during the 2 s period of interest preceding a response were then divided by mean log transformed power values in each frequency band a baseline period corresponding to the final 0.2 s preceding stimulus onset. Data from time windows either side of the period of interest were discarded after convolution to avoid edge effects, before averaging across trials (**Figure 1C)**.

##### Source Reconstruction

MEG source localisation was conducted using the linearly constrained minimum variance (LCMV) beamformer in SPM12 with a single-shell forward model to generate maps of the mean source power difference between conditions on a 10 mm grid^49^, co-registered to MNI coordinates. Theta (4-8 Hz) power was estimated in the 1.4 to 0.6s preceding response for each trial. The frequency bandwidth and time window were selected based on the sensor level time-frequency results **(Figure 1C**). The summary statistic computed for changes in theta power was then averaged across trials for each condition. Summary images for each participant were entered into a second level paired-sample t-test in SPM12, comparing naming versus control conditions, with a voxel-level family-wise error (FWE) corrected significance threshold of *P* < 0.05 and a cluster extent of at least 10 voxels. All image coordinates are listed in Montreal Neurological Institute (MNI) space (Figure 1D). Voxels that survived this significance threshold were then used as seeds for the theta phase coupling analysis.

#### Theta Phase Coupling

Instantaneous theta phase in the 1s preceding response onset in each voxel across the brain was obtained by applying the Hilbert transform to the 4-8 Hz band-pass filtered time series generated by the LCMV beamformer. The 1s time window was selected to account for the mean RT of both naming and control conditions – both of which showed a normal distribution around 1s – and remain suitable for at least 4 theta cycles. The left lingual and left mPFC seed voxel for each participant was selected as that with the greatest theta power increase in naming > control within 20 mm of the group maximum co-ordinates [left lingual gyrus MNI co-ordinates (-14; -96; -12); left mPFC MNI (-8; 56; -10)] (**Figure 2A**) ^37^. The group maximum co-ordinates were obtained as described above (see ‘Source Reconstruction’). Theta phase coupling was then estimated between a single seed voxel and every other voxel in the brain using the Phase Locking Value (PLV) ^50^. The PLV is computed as the resultant vector length of phase differences between source and seed voxels over time, such that a larger value indicates less variance in the phase difference between two signals. Summary images for each participant – containing trial averaged PLV values in each voxel across the whole brain for each condition – were then entered into a second level paired-sample t-test in SPM12 contrasting naming > control, with a voxel-level significance threshold of *P* < 0.001 and cluster-level significance threshold of FWE corrected *P* < 0.05. Although PLV measures phase coupling, it can be biased by concurrent changes in power that affect the signal-to-noise ratio^51^. To eliminate this potential confound, we used linear regression to remove any effect of trial-by-trial variance in mean-corrected power across naming and control trials and then contrasted the residual PLV values^37,52^.

Finally, to assess whether the spatial overlap between the two phase-coupled networks exceeded chance, we quantified voxel-wise overlap between the left lingual-seeded theta phase-coupled network and the left mPFC-seeded theta phase-coupled network. Both networks were defined as binary significance maps derived from second-level SPM analyses and constrained to the same analysis mask. The expected overlap under the null hypothesis of spatial independence was computed as *E*= (*n*_l_ x *n*_2_)/*N*, where *n*_l_ and *n*_2_ are the numbers of suprathreshold voxels in each network and N is the number of voxels in the SPM analysis mask. Statistical significance was assessed using a permutation test (10,000 iterations), in which voxel labels were randomly reassigned within the analysis mask while preserving cluster sizes, generating a null distribution of overlap values (Supplementary Figure 1).

#### Phase-Amplitude Coupling

Theta-gamma phase-amplitude coupling (PAC) was estimated across the whole brain volume in the 1s preceding response. In line with the theta phase coupling analysis, the 1s time window was selected to account for the mean RT of both naming and control conditions and remain suitable for at least 4 cycles of theta. Instantaneous gamma (40 – 100 Hz) amplitude and theta (4-8 Hz) phase for each voxel were obtained by applying the Hilbert transform to source reconstructed time series. Theta-gamma PAC was then estimated independently in each voxel using the mean vector length (MVL)^16^. The MVL is computed as the resultant vector length of theta phase values in each sample, weighted by the concurrent gamma amplitude in the same sample, such that larger values indicate greater phase amplitude coupling. Summary images for each subject were then entered into a second level paired-samples t-test in SPM12, contrasting naming > control conditions. We used a significance threshold of *P* < 0.001 at the voxel-level and FEW-corrected *P* < 0.05 at the cluster-level. (**Figure 3A)**. To ensure our results were not biased by concurrent changes in power that can affect the signal-to-noise ratio ^51^, we used linear regression to remove any effect of trial-by-trial variance of both theta and gamma power, then used a paired samples t-test to compare the residual MVL values between conditions (naming versus control) in any regions of interest (ROIs) that showed significant PAC ^37,52^.

Next, we used a comodulogram analysis to identify the individual theta and gamma frequency pairs that showed significant PAC during naming compared to control ^16^ (**Figure 3B**). Using the LCMV beamformer algorithm, timeseries were obtained from two 20mm radius seeds centred around each of the voxels that showed the greatest theta-gamma PAC during naming vs control in the whole-brain analysis described above. These voxels were in the left pars triangularis of the inferior frontal gyrus (LIFG, MNI [-38; 28; 2]) and left fusiform area (MNI [-28; -10; -30]). Using source reconstructed time series, MVL was computed across theta phases in the 4-8 Hz band (in 1 Hz steps) and gamma amplitudes in the 40-100 Hz band (in 2 Hz steps), for the 1s period preceding response onset in each trial ^53^. These values were then averaged to obtain a single MVL modulation index (MI) value per amplitude and phase, or phase-amplitude matrix, for each condition (the “comodulogram”)^16^. Comodulograms for each condition were contrasted using non-parametric cluster-based statistics, which has been shown to adequately control the type-I error rate for electrophysiological data^54^. First, an uncorrected paired-samples t-test was performed (naming versus control), and all MVL-MI values exceeding a 5% significance threshold were grouped into clusters. The maximum t-value within each cluster was carried forward. Next, a null distribution was obtained by randomizing the condition label (naming/control) 10,000 times and calculating the largest cluster-level t-value for each permutation. The maximum t-value within each original cluster was then compared against this null distribution, with values exceeding a threshold of p < 0.025 deemed significant.

Finally, to assess whether trial-by-trial fluctuations in theta-gamma PAC related to behavioural performance, linear mixed-effects models were fitted to single-trial RTs using the lme4 package in R (v4.5.2; R Core Team, 2025). Within each participant, PAC values were z-scored to isolate trial-to-trial deviations from each individual’s mean coupling strength. Models included PAC strength, task condition (clear naming vs noise control), and their interaction as fixed effects, with random intercepts for participant; follow-up analyses additionally included random slopes for condition to assess robustness to inter-individual variability in task effects. Separate models were fitted for left inferior frontal gyrus and fusiform cortex regions of interest. Statistical inference was based on Satterthwaite-approximated *t*-tests, with Type III ANOVA used to evaluate interaction effects.

---

**Box 1.**
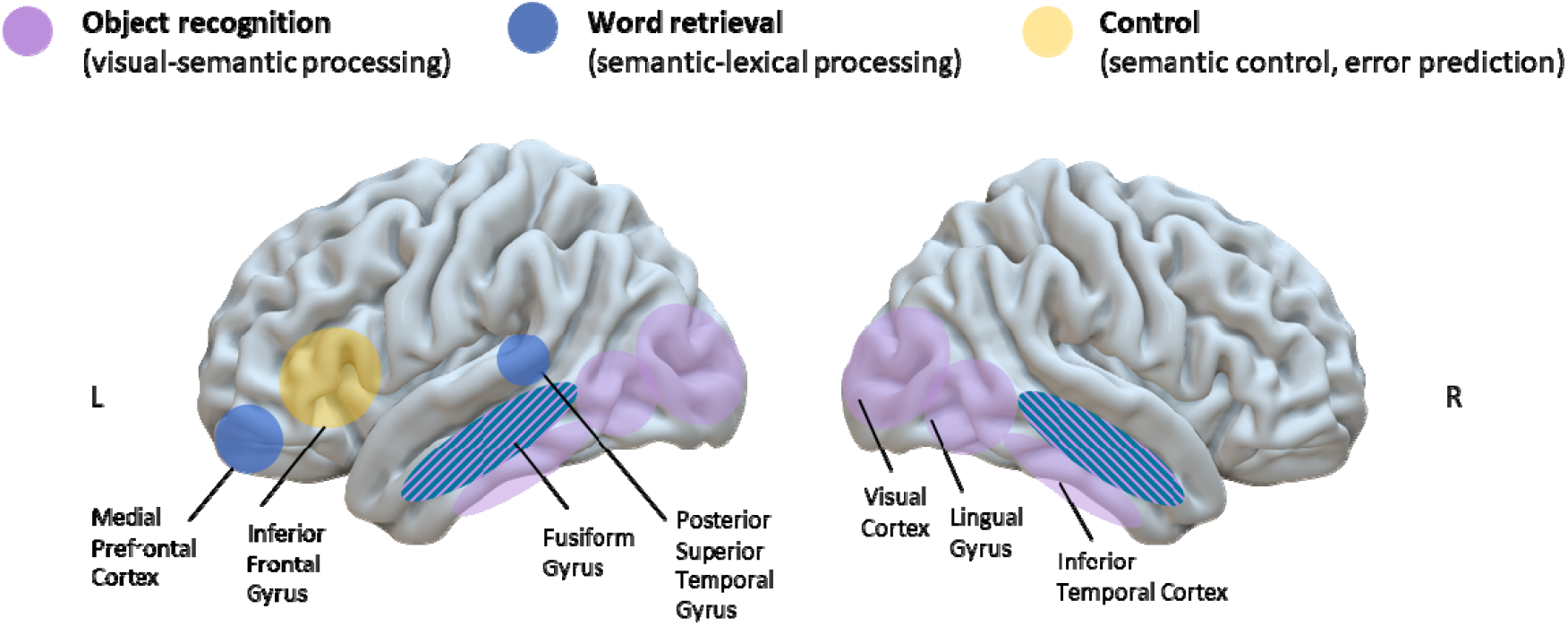
Pre-articulatory language production network during picture naming. Picture naming engages a distributed network supporting object recognition, semantic-lexical retrieval and control prior to articulation. Visual-semantic processing along the ventral visual stream (lingual, fusiform, and inferior temporal cortex) supports object recognition. Medial prefrontal and medial temporal regions contribute to word retrieval, encompassing semantic-lexical processes (word meaning and form), while the posterior superior temporal gyrus supports phonological processing of the retrieved word. The inferior frontal cortex mediates semantic control and selection among competing lexical representations. Frontal contributions are predominantly left-lateralised, whereas temporal regions show relatively bilateral involvement. The fusiform gyrus (striped) participates in both conceptual and lexical stages, with modality-dependent lateralisation, and is preferentially right-lateralised during visually guided word retrieval.

## Notes

### Competing Interest Statement

The authors have declared no competing interest.

### Summary of Updates

Paper updated to include supplemental file.

