## Supplemental Material for "Theta phase and theta-gamma coupling organise the spoken language network"

| Anatomical Location | Extent (voxels) | Z | MNI co-ordinates | | |
| --- | --- | --- | --- | --- | --- |
|  |  |  | x | y | z |
| **Left Lingual Gyrus** | **14646** | **6.35** | **-14** | **-96** | **-12** |
| Right Lingual Gyrus |  | 6.26 | 14 | -94 | -16 |
| Left Calcarine Cortex |  | 6.18 | -8 | -96 | -12 |
| Left Lingual Gyrus |  | 6.00 | -14 | -88 | -8 |
| Left Lingual Gyrus |  | 5.80 | -24 | -96 | -14 |
| Right Lingual Gyrus |  | 5.65 | 22 | -96 | -14 |
| Left Calcarine Cortex |  | 5.52 | 2 | -98 | -8 |
| Left Superior Occipital Cortex |  | 5.52 | -14 | -96 | 6 |
| **Left Medial Pre-Frontal Cortex** | **697** | **5.29** | **-8** | **56** | **-10** |
| Left Medial Pre-Frontal Cortex |  | 5.29 | -10 | 62 | -8 |

### **Supplementary Table 2A**

### Peak cortical source estimates of theta (4-8 Hz) **power** increases in naming versus control

### **Supplementary Table 1**

| Anatomical Location | Extent (voxels) | Z | MNI co-ordinates | | |
| --- | --- | --- | --- | --- | --- |
|  |  |  | x | y | z |
| **Right Superior Occipital Gyrus** | **3994** | **4.41** | **26** | **-74** | **12** |
| Right Lingual Gyrus |  | 4.14 | 26 | -56 | -6 |
| Right Middle Fusiform Gyrus |  | 3.93 | 24 | -48 | -14 |
| Right Lingual Gyrus |  | 3.86 | 18 | -54 | -6 |
| Right Lingual Gyrus |  | 3.85 | 26 | -62 | -4 |
| Right Posterior Fusiform Gyrus |  | 3.75 | 40 | -50 | -24 |
| Right Middle Fusiform Gyrus |  | 3.66 | 32 | -48 | -12 |
| Right Posterior Fusiform Gyrus |  | 3.64 | 30 | -74 | -2 |

### Peak cortical sources of theta (4–8 Hz) **phase coupling** with the **left lingual gyrus seed** [MNI (-14; -96; -12)] during naming versus control.

| Anatomical Location | Extent (voxels) | Z | MNI co-ordinates | | |
| --- | --- | --- | --- | --- | --- |
|  |  |  | x | y | z |
| **Right Parahippocampal Gyrus** | **6061** | **4.75** | **20** | **-32** | **-14** |
| Right Anterior Lingual Gyrus |  | 4.44 | 18 | -44 | -10 |
| Right Fusiform Gyrus |  | 4.42 | 20 | -43 | -11 |
| Left Precuneus |  | 4.01 | -16 | -48 | 2 |
| Right Hippocampus |  | 3.71 | 34 | -10 | -14 |
| Right Posterior Cingulate Cortex |  | 3.36 | 4 | -38 | 8 |

### **Supplementary Table 2B**

### Peak cortical sources of theta (4–8 Hz) **phase coupling** with the **left medial pre-frontal cortex seed** [MNI (-8; 56; -10)] during naming versus control.

| Anatomical Location | | Extent (voxels) | | Z | MNI co-ordinates | | |
| --- | --- | --- | --- | --- | --- | --- | --- |
|  |  |  |  |  | x | y | z |
| **Left Inferior Frontal Gyrus (Pars Triangularis)** | | **405** | | **4.45** | **-38** | **28** | **2** |
| Left Inferior Frontal Gyrus (Pars Orbitalis) | |  | | 4.26 | -42 | 34 | -4 |
| Left Inferior Frontal Gyrus (Pars Orbitalis) | |  | | 3.23 | -46 | 28 | -12 |
| **Left Fusiform** | | **352** | | **4.25** | **-28** | **-10** | **-30** |
| Left Inferior Temporal Gyrus | |  | | 3.52 | -38 | -22 | -22 |
|  | Cortical sources of theta (4–8 Hz) gamma (40–100 Hz) **phase-amplitude coupling** during naming versus control. | |  | |  |  |  |

### **Supplementary Table 3**

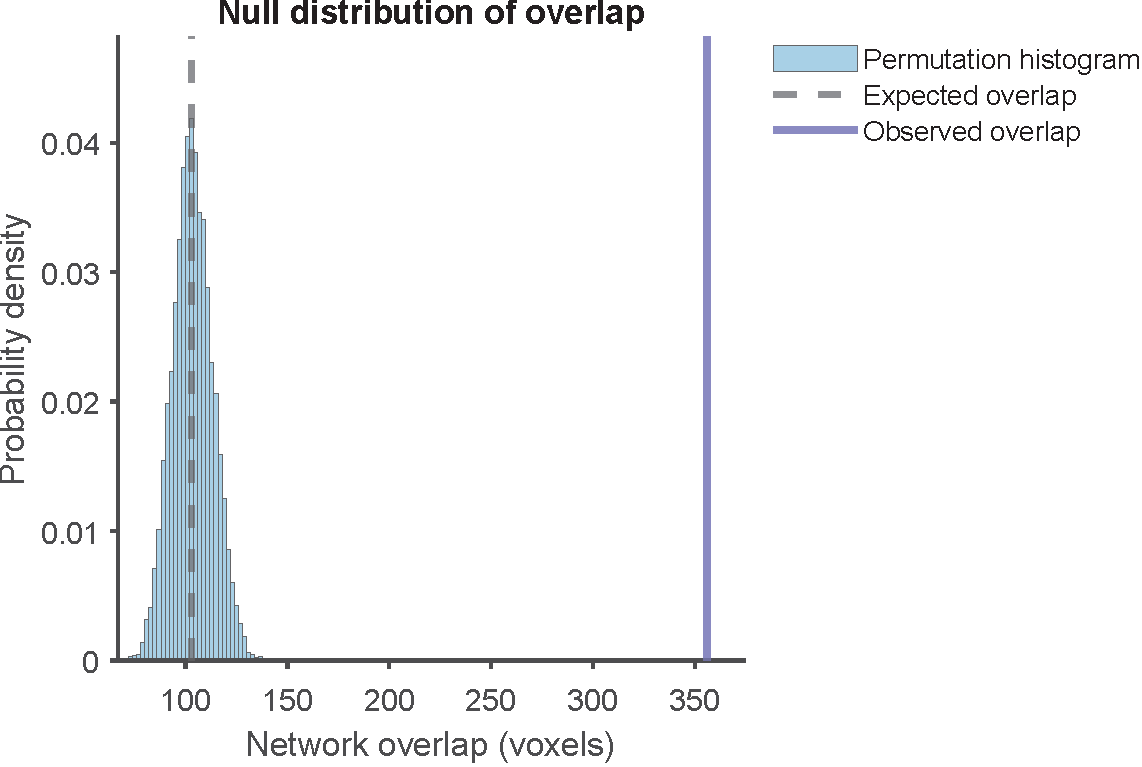

### **Supplementary Figure 1. Spatial overlap between theta phase-coupled networks exceeds chance expectations.** Null distribution of voxel-wise overlap between the left lingual-seeded and medial prefrontal cortex (mPFC)-seeded theta (4-8 Hz) phase-coupling networks (see main-text Figure 2). Blue bars show the permutation-derived null distribution of overlap values obtained by randomly reassigning voxel labels within the second-level SPM analysis mask while preserving network sizes (10,000 permutations; see STAR Methods). The dashed grey line indicates the expected overlap (103.1 voxels), and the solid purple line marks the observed overlap between networks (356 voxels). The observed overlap significantly exceeded chance expectation (p < 0.001), indicating non-random spatial convergence between ventral occipito-temporal and medial fronto-temporal theta coherence networks.

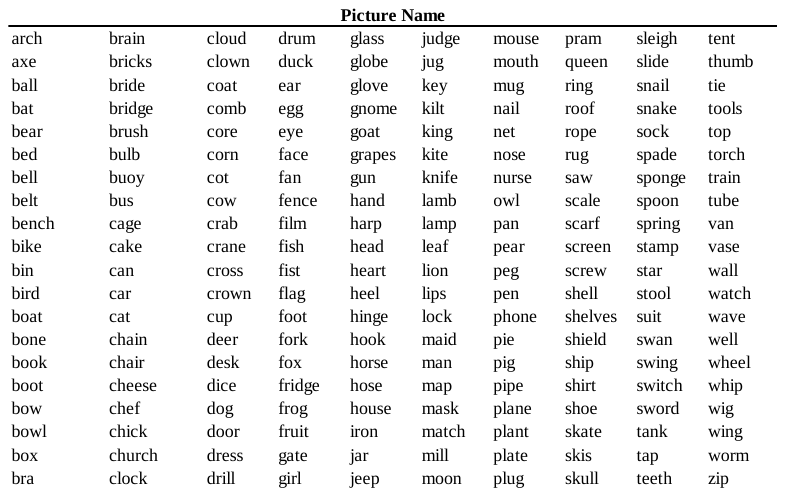

### List of the target words for common objects used in the naming condition of the picture naming task. Target words drawn from the International Picture Naming Project (IPNP; http://crl.ucsd.edu/experiments/ipnp/index.html; Szekely et al., 2004). All target words are monosyllabic with high naming-agreement.

### **Supplementary Table 4**
